# Sequence-only antigen context improves modeling of antibody hypervariable regions

**DOI:** 10.64898/2026.10.19.760314

**Authors:** Faisal Bin Ashraf, Stefano Lonardi

## Abstract

The complementarity-determining regions (CDRs) of antibodies govern antigen specificity; consequently, CDR design should be performed conditional on the target antigen. Most published approaches depend on the availability of the antigen’s three-dimensional (3D) structure, which is often difficult to obtain experimentally. In contrast, sequence-only models circumvent this requirement but are typically trained on antibodies without incorporating antigen context; exposure to antigen sequences alone does not necessarily entail learning the underlying interaction determinants. Here, we introduce LangAAI, a masked language model that conditions exclusively on antibody and antigen primary sequences and therefore does not require 3D structures, docking poses, or epitope annotations. To assess whether the model exhibits genuine antigen conditioning, we replace the cognate antigen with a non-binding decoy. Whereas antigen-agnostic models are largely insensitive to this substitution, LangAAI shows a marked reduction in sequence recovery that is specifically concentrated in the hypervariable CDR3 loop. Across benchmarks, LangAAI improves amino-acid recovery relative to existing antibody language models, matches the performance of structure-based methods when evaluated under their respective protocols, and ranks experimentally measured variants by affinity comparably to inverse-folding models that use backbone coordinates. Collectively, these results support LangAAI as a structure-independent approach for antigen-aware antibody optimization, enabling prioritization of candidate variants for targets lacking solved antigen–antibody complexes.

## 1. Introduction

About two-thirds to three-quarters of any antibody consists of constant amino acid sequences. An antibody recognizes its target (i.e., the *epitope* of an antigen) through six *complementarity-determining regions* (CDRs), three of which are located on the heavy chain (H1, H2, H3) and three on the light chain (L1, L2, L3). The specificity of the CDR-epitope recognition is the foundation of humoral immunity, diagnostics, and therapeutic design [Carter and Rajpal, 2022]. Computationally designing CDRs for a specific antigen requires one to solve the inverse of the molecular recognition problem, but the combinatorial space is too large to search. For instance, the length of CDR-H3 is about 15 amino acids [Briney et al., 2019], thus there are 20^15^ candidate sequences for CDR-H3. This vastly outnumbers the 10^10^–10^12^ variants achievable in physical display libraries [Bradbury et al., 2011]. Computational design of antibody seeks to limit the search space *in silico*, but current computational approaches face two critical limitations. First, they require the knowledge of the 3D structure for the antigen. Yet, fewer than 8,000 antibody–antigen complexes exist in the Protein Data Bank [Berman et al., 2000; Dunbar et al., 2014] and state-of-the-art docking algorithms frequently fail at predicting the structures of these complexes because these interactions lack a shared evolutionary history and involve highly flexible hypervariable binding regions. For instance, AlphaFold-Multimer and AlphaFold3 [Abramson et al., 2024] achieve accurate predictions in only 11% [Yin et al., 2022] and 8.9% [Hitawala and Gray, 2024] of cases, respectively. Second, while the primary sequence of antibodies is abundant (e.g., two billion entries in the Observed Antibody Space [Olsen et al., 2022]), it is highly skewed toward a small number of intensively studied targets (e.g., the SAR-Cov22 spike protein). A sequence model trained on this highly biased corpus can minimize reconstruction error merely by memorizing antibody statistics, entirely ignoring the target antigen. Therefore, any sequence-based model must explicitly address the role of the antigen to understand and generate CDRs.

Antibody discovery increasingly formulates CDR design as a machine learning problem: given an antigen, propose a complementarity-determining region likely to bind it. Three largely disjoint communities have approached some part of this task, yet none has achieved a sequence-only generative solution that generalizes to unseen targets. First, the antibody language model community has produced strong protein language models fine-tuned on antibody repertoires, including AntiBERTy [Ruffolo et al., 2021], AbLang2 [Olsen et al., 2024], and IgLM/IgBert [Shuai et al., 2023]. However, these are trained on the antibody chain in isolation, while the target antigen never enters the model. Consequently, the model proposes identical CDRs irrespective of the intended target. Second, structure-based CDR design methodologies—such as MEAN [Kong et al., 2022], dyMEAN [Kong et al., 2023], DiffAb [Luo et al., 2022], RAAD [Wu et al., 2025], RADAb [Wang et al., 2025], AbFlowNet [Abir et al., 2025], AbMEGD [Chen et al., 2025], and dyAb [Tan et al., 2025] do condition on the antigen. Yet, they rely strictly on resolved or predicted 3D coordinates and explicit epitope-contact definitions, which are often difficult to obtain experimentally. A third family comprises general protein–protein interaction language models, such as MINT [Ullanat et al., 2026], PPLM [Liu et al., 2026], PLM-interact [Liu et al., 2025]. While these methods process both sequences without structural priors, they lack antibody-specific awareness and are trained strictly to discriminate interacting pairs rather than to generatively reconstruct binding loops. As we demonstrate below, merely providing access to the antigen sequence does not inherently yield true antigen-conditioned generation unless the architecture is explicitly trained to learn from two sequences within an antibody–antigen context. To the best of our knowledge, no pure-sequence antibody language model has been trained to be genuinely antigen-conditioned across diverse targets. The closest existing approach, PALM-H3 [He et al., 2024], pairs an ESM-2 antigen encoder with an antibody decoder via cross-attention, but it was developed and evaluated exclusively on SARS-CoV-2 antigens, providing no evidence of transferability across distinct antigen families. Ultimately, what has been missing is both a model that conditions on the antigen sequence alone and a rigorous evaluation capable of demonstrating that this conditioning is functionally authentic.

In response to these needs, here we present LangAAI, an antigen-conditioned masked language model that processes antibody–antigen sequence pairs via a co-attention-based joint transformer architecture. The model is trained on paired sequences alone, without coordinates, docking poses, or epitope definitions. To the best of our knowledge LangAAI is the first general-purpose language model of joint antibody–antigen sequences. Because baseline reconstruction accuracy cannot differentiate true antigen conditioning from the mere memorization of antibody sequence priors, we introduce a paired-shuffle metric contrasting cognate antigens against decoys. Across held-out evaluation sets, LangAAI consistently outperforms established language model baselines in sequence recovery, particularly within the critical CDR-H3 loop. Crucially, replacing the true antigen significantly degrades predictive accuracy, an effect that our ablation tests trace directly to the CDRs rather than the constant framework. Furthermore, when assessed on external structural benchmarks, LangAAI rivals explicitly structure-conditioned methods, indicating that paired sequences implicitly capture the biophysical context normally derived from spatial geometry. Finally we demonstrate two key capabilities of LangAAI’s sequence probabilities. First, they accurately rank experimental variants by measured binding affinity, outperforming all sequence-only baselines and matching the performance of leading structure-based models. Second, using these probabilities to generate new candidate sequences yields a significantly higher fraction of improved binders compared to the standard experimental campaigns. In summary, these results suggest that antigen-conditioned design does not absolutely require explicit 3D structure, but rather an objective that makes the antigen an active determinant.

## 2. Results

### 2.1. Overview of LangAAI

LangAAI is an antigen-conditioned masked language model, designed so that the reconstruction of any masked antibody residue strictly depends on its paired antigen. Figure 1a shows an overview of LangAAI. First, a frozen protein language model produce independent embedding of both the antibody and antigen sequences. Each embedding then passes through a dedicated, lightweight trainable adapter, which projects both representations into a shared latent space. By freezing the underlying language model, we restrict parameter optimization exclusively to the interaction module. Next, the adapted representations are concatenated alongside a prepended `[CLS]` token, with segment identifiers used to distinguish the respective sequences. A stack of transformer layers process this joint sequence, enabling direct co-attention between antibody and antigen residues. This co-attention mechanism provides the exclusive pathway for antigen context to influence antibody reconstruction, and so any performance gains relative to an antigen-agnostic baseline can be robustly attributed to the specific antigen pairing rather than to intrinsic antibody sequence statistics.

**Figure 1:**
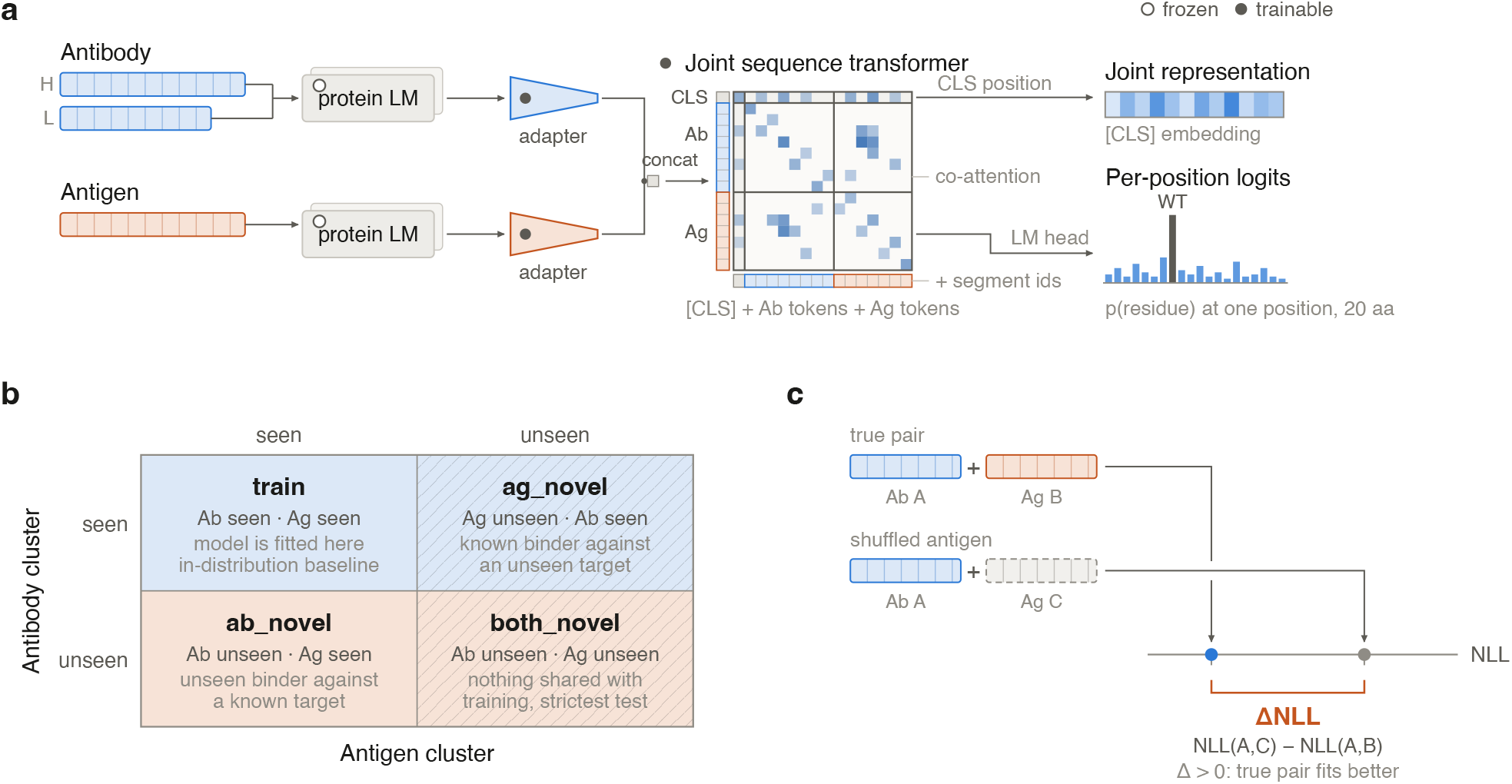
LangAAI model overview, entity-holdout evaluation, and paired-shuffle specificity metric. **(a)** The sequence of antibody and antigen are processed through frozen protein language models, adapters, and a co-attention Joint Sequence Transformer to generate per-position binding logits. **(b)** Entity-holdout splits partition data by sequence clusters to evaluate model generalization to unseen entities. **(c)** The Paired-Shuffle Delta Metric assesses pair prediction accuracy using Δ*NLL* = *NLL*(*A, C*) *− NLL*(*A, B*) to contrast true versus shuffled antigens.

The joint transformer stack yields two complementary readouts. First, a language-model head generates a probability distribution over the 20 canonical amino acids at each masked antibody position. We utilize this distribution as the primary training objective (Figure 1a, right). Second, the `[CLS]` token embedding yields a fixed-length joint representation of the molecular complex. Our masking strategy selectively targets the complementarity-determining regions (CDRs) and the CDR-flanking regions with a specific emphasis on CDR3. We avoid masking framework positions, as their sequences are highly conserved and recoverable from baseline germline context. Moreover, doing so would allow the model to minimize the loss function without considering the antigen at all.

Because sequence similarities across both antibodies and antigens acts as strong confounders, we evaluated the model under a two-dimensional split rather than a single random partition (Figure 1b). We first clustered antibody and antigen sequences independently based on their respective sequence similarities. Upon pairing these molecules, their assigned clusters (seen versus unseen) established four distinct evaluation regimes. In the `train` regime, both the antibody and antigen clusters are present in the training data. The `ag_novel` regime evaluates a previously observed antibody against a novel antigen cluster. The `ab_novel` regime pairs an unseen antibody with a known antigen target. Finally, in the `both_novel` regime, neither cluster was observed during training. The `train` partition inherently serves as both the training corpus and the in-distribution baseline. Conversely, the `both_-novel` regime provides the most rigorous assessment of model generalization, strictly precluding the network from relying on memorized antibody–antigen interactions.

To evaluate antigen-conditioning directly, we implemented a paired shuffling protocol (Figure 1c). For a held-out antibody *A* and its true cognate antigen *B*, we calculated the negative log-likelihood (NLL) of the masked antibody residues. We then computed the NLL for a shuffled complex by replacing *B* with an unrelated decoy antigen *C*, sampled from the identical evaluation regime. By calculating the difference, ΔNLL = NLL(*A, C*) *−* NLL(*A, B*), we established a comparative metric wherein a positive ΔNLL indicates that the true antigen provides a superior structural explanation for the antibody sequence compared to an arbitrary target. Because the antibody, the masking pattern, and the model are identical across the two calculations, this contrast isolates the contribution of antigen identity and cannot be satisfied by a model that has learned antibody sequence priors alone.

### 2.2. Antigen conditioning improves CDR reconstruction and generalizes to novel antibody-antigen

We evaluated the masked sequence reconstruction of antibody complementarity-determining regions (CDRs) by comparing LangAAI against five existing language models. Our benchmark utilizes 17, 983 held-out antibody–antigen pairs separated into three generalization regimes: `ab_novel` (*n* = 8, 551), `ag_novel` (*n* = 8, 804), and `both_novel` (*n* = 628). The five baselines span antibody-specific models (AbLang2 [Olsen et al., 2024], AntiBERTy [Ruffolo et al., 2021]), a general protein model (ESM-2 [Lin et al., 2023]), and protein–protein interaction networks (MINT [Ullanat et al., 2026], PLM-interact [Liu et al., 2025]). While LangAAI, MINT, and PLM-interact utilize the paired antigen sequence as input, the other three models score the antibody independently.

Although the data split strictly segregates clusters based on overall sequence homology, we determined that 66.6% of the held-out set’s CDR-H3 sequences overlap with the training corpus (see Methods for details). Consequently, we report every benchmark comparison independently for complexes whose CDR-H3 was exposed during training and those whose CDR-H3 was entirely unseen. We first quantified whole-sequence recovery by pooling predictions across every scored residue, encompassing both the flanking and CDR1–3 regions. Figure 2a shows that LangAAI achieved the highest recovery rate in five of the six evaluation cohorts, accounting for both complex novelty and CDR-H3 overlap. For sequences containing a previously seen CDR3, LangAAI outperformed the strongest baseline across all three novelty regimes: `ab_novel` (0.842 versus 0.674), `ag_novel` (0.912 versus 0.619), and `both_novel` (0.882 versus 0.694). When evaluated on unseen CDR3 sequences, LangAAI maintained its advantage in the `ab_novel` (0.687 versus 0.619) and `both_novel` (0.734 versus 0.594) cohorts. The single exception occurred within the `ag_novel` regime. Here, LangAAI performed comparably to the leading baselines, AntiBERTy and AbLang2 (0.645 versus 0.658 for both), exhibiting no statistically significant deficit.

**Figure 2:**
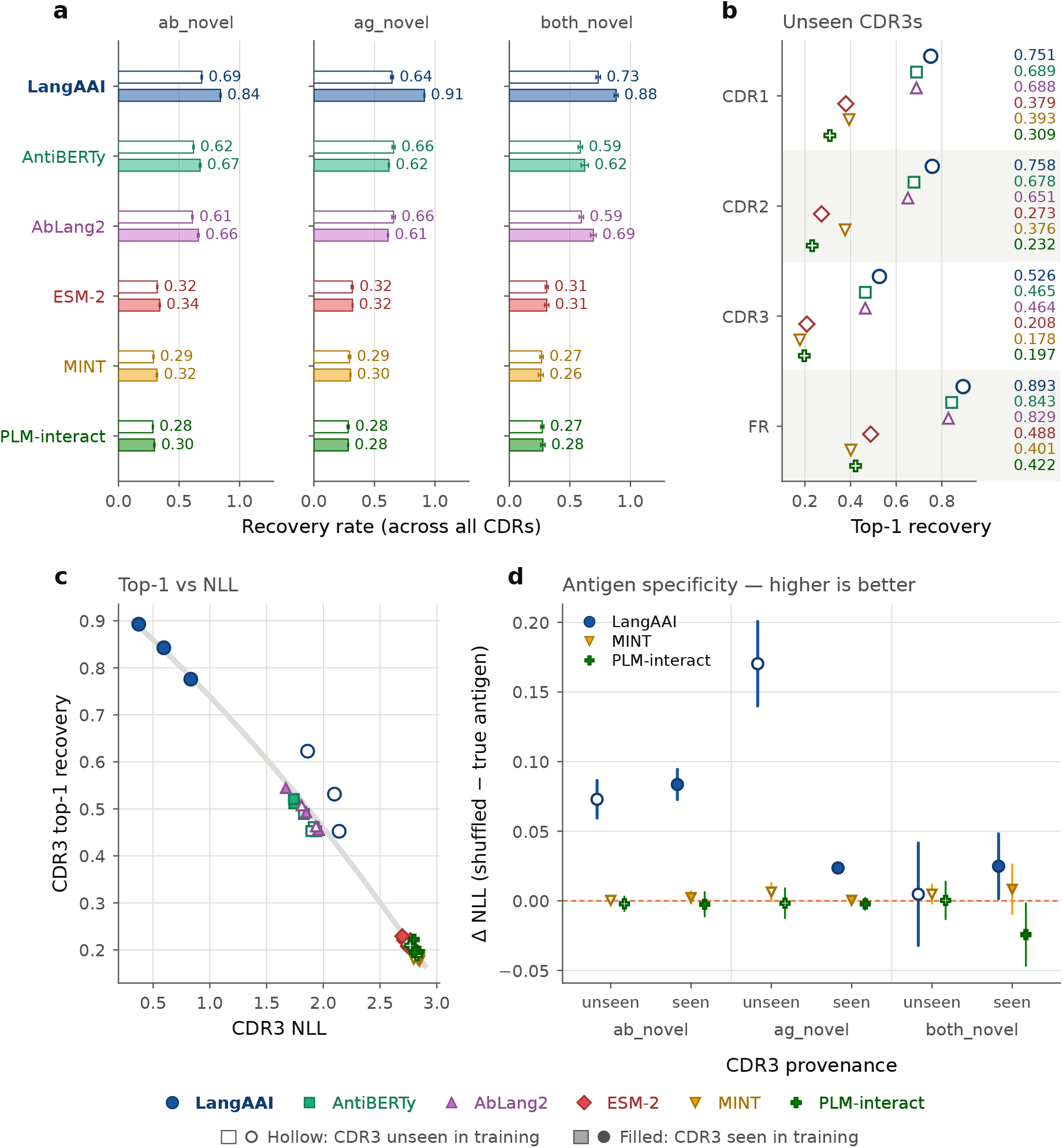
Amino Acid Reconstruction, CDR3 provenance, and antigen specificity. **(a)** Whole-sequence top-1 recovery, split by CDR3 provenance within each novelty regime; bars are group means and error bars are 95% confidence intervals over examples. **(b** Top-1 residue recovery by region for all six models, restricted to examples with a unseen CDR3 and pooled across novelty regimes. **(c)** Mean CDR3 NLL against mean top-1 recovery, one point per model *×* regime *×* provenance cell; all models trace a shared accuracy–NLL relation (grey); LangAAI’s seen-CDR3 cells sit at the favourable extreme of that same relation, whereas its unseen-CDR3 cells sit above it. **(d)** Antigen specificity, measured as Δ*NLL* = *NLL*(shuffled antigen) *−NLL*(true antigen); higher is better, and Δ = 0 (dashed line) means antigen identity carries no information for the model; points are means with 95% confidence intervals.

To evaluate performance in the absence of memorization, we restricted our analysis to unseen CDR-H3 sequences. Under this stringent condition, LangAAI recovered masked residues more accurately than the top-performing baseline (AntiBERTy) across all four structural domains: CDR1 (0.751 vs. 0.689), CDR2 (0.758 vs. 0.678), CDR3 (0.526 vs. 0.465), and flanking regions (0.893 vs. 0.843) (see Figure 2b). All architectures exhibited an identical performance pattern — framework residues were the easiest to reconstruct and CDR3 the hardest — reflecting the extreme sequence diversity of CDR-H3. Notably, while the antibody-specific baselines strongly outperformed ESM-2, the generalized protein interaction models failed to exceed it.

We observed a robust anti-correlation between top-1 sequence recovery and negative log-likelihood (NLL) across all architectures (*r* = *−*0.89 for CDR3), conforming to a shared predictive relationship (Figure 2c). Rather than deviating from this universal relationship, which would strongly indicate simple sequence memorization, LangAAI’s seen CDR-H3 test samples map precisely to its most favorable extreme. More importantly, the model’s unseen CDR-H3 test samples deviate positively, positioning strictly above the standard curve. This upward shift demonstrates that for novel sequences, LangAAI yields significantly more accurate top-1 predictions than its corresponding likelihood scores would predict. Combined with LangAAI’s established dominance on unseen sequences across all regional domains (Figure 2b) and within two of the three whole-sequence recovery regimes (Figure 2a), these properties confirm that the LangAAI’s efficacy extends beyond exposure bias to enable the accurate reconstruction of genuinely novel CDR-H3 targets.

### 2.3. LangAAI leverages antigen context to reconstruct novel antibodies

It is obvious that reconstruction accuracy is insufficient to confirm whether the model explicitly conditions on the paired antigen. To isolate this latter variable, we define the *differential negative log-likelihood* (ΔNLL) by comparing the reconstruction error of the true complex against a decoy pair featuring a shuffled antigen partner. Formally, ΔNLL = NLL(shuffled) *−* NLL(true). A positive ΔNLL value indicates that the model utilizes the antigen identity to improve sequence reconstruction over an arbitrary pairing. A value of ΔNLL = 0 implies a complete absence of informative antigen conditioning. The three antibody-specific baselines, which lack structural antigen awareness and thus intrinsically produce ΔNLL = 0, were omitted from this evaluation.

Figure 2d shows that, among the three antigen-conditioned models, LangAAI produced the largest ΔNLL in five of the six regime by cdr3-exposure, and was the only model with any interval excluding zero in the positive direction. MINT’s 95% CIs encompassed zero in all six partitions, and PLM-interact’s CIs included zero in five, with the remaining subset (`both_novel`, seen CDR3) deviating negatively. LangAAI had the clearest structural specificity within the `ab_novel` regime for both unseen (ΔNLL = 0.073, 95% CI 0.059–0.087) and seen (ΔNLL = 0.084, 95% CI 0.073–0.094) CDR3 targets. The highest antigen dependence was observed within the `ag_novel` regime exclusively on unseen CDR3s (ΔNLL = 0.170, 95% CI 0.140–0.201; *t* = 11.0, *P* = 7.3 *×* 10^*−*26^), a 7.2-fold increase over the corresponding seen CDR3 cohort in the same regime (ΔNLL = 0.024; Welch *t* = 9.4, *P* = 7.8 *×* 10^*−*20^).

Because the model’s antigen dependency is strongest where structural memorization is impossible, this indicates that LangAAI explicitly recruits binding-partner data to reconstruct unfamiliar antibody targets.

We also observed two additional analytical qualifications. First, the specificity is statistically indistinguishable from zero for unseen CDR3s in the underpowered `both_novel` regime (ΔNLL = 0.005, 95% CI *−*0.032 to 0.042). Second, the observed effect (ΔNLL = 0.170) represents a highly specific but magnitude-limited perplexity ratio of 1.19. To ensure this signal is not an artifact of data leakage, we evaluated the impact of near-duplicate sequences. While 32.4% of unseen examples are within two edits of a training sequence, excluding these near-matches from the unseen `ag_novel` subset increased the differential log-likelihood (ΔNLL = 0.194, 95% CI 0.157–0.231). Thus, antigen specificity is structurally robust and not a byproduct of data leakage due to sequence homology.

### 2.4. Sequence-only design with LangAAI is competitive with structure-based methods

CDR design is dominated by methods that condition on the 3D structure of antibody–antigen complex. To position LangAAI against that mainstream without retraining any baseline, we evaluated it on CHIMERA-Bench [Ahmed et al., 2026]. This standardized benchmark provides a fixed 292-complex `epitope_group` test split and reports per-CDR Amino Acid Recovery (AAR) for up to ten structure-based methods, all uniformly retrained by the benchmark authors under identical conditions. These baselines encompass equivariant GNNs (RAAD [Wu et al., 2025], MEAN [Kong et al., 2022], dyMEAN [Kong et al., 2023]), diffusion models and their extensions (DiffAb [Luo et al., 2022], AbMEGD [Chen et al., 2025], RADAb [Wang et al., 2025], AbFlowNet [Abir et al., 2025]), flow matching (dyAb [Tan et al., 2025]), autoregressive generation (RefineGNN [Jin et al., 2021]), hierarchical equivariant networks (AbDockGen [Jin et al., 2022]), and conjoined ODEs (AbODE [Verma et al., 2023]). To ensure a rigorous comparison for the redesign-from-scratch task executed by these models, we masked each CDR loop in its entirety and utilized CHIMERA-Bench’s unmodified aar() function and defined CDR-span resolutions. Importantly, whereas the baseline architectures depend on 3D spatial data, LangAAI operates exclusively on the raw antigen sequence, receiving no coordinate inputs or predefined epitope boundaries.

Figure 3a shows that, when evaluated under this matched protocol, LangAAI achieved the highest Amino Acid Recovery (AAR) among all evaluated methods in four of the six CDR regions, and the second-highest AAR in the remaining two. Comparing LangAAI directly to the strongest region-specific baseline, LangAAI yielded competitive or superior recovery in CDR-H1 (0.731 vs. RAAD 0.72), CDR-H2 (0.67 vs. RAAD 0.67), CDR-H3 (0.422 vs. MEAN 0.42), CDR-L1 (0.627 vs. RADAB 0.64), CDR-L2 (0.703 vs. AbFlowNet 0.51), and CDR-L3 (0.620 vs. DiffAb 0.43). However, only five of the ten leaderboard methods report light-chain CDRs, and the four strongest heavy-chain performers are not among them. So, LangAAI’s large margins in CDR-L2 and CDR-L3 are measured against a comparatively weaker field. In addition, the average loop length varies more than four-fold across regions (CDR-L2 3.1 residues, CDR-H3 15.5). With AAR rising as loops shorten for every method, only within-region comparisons are meaningful.

**Figure 3:**
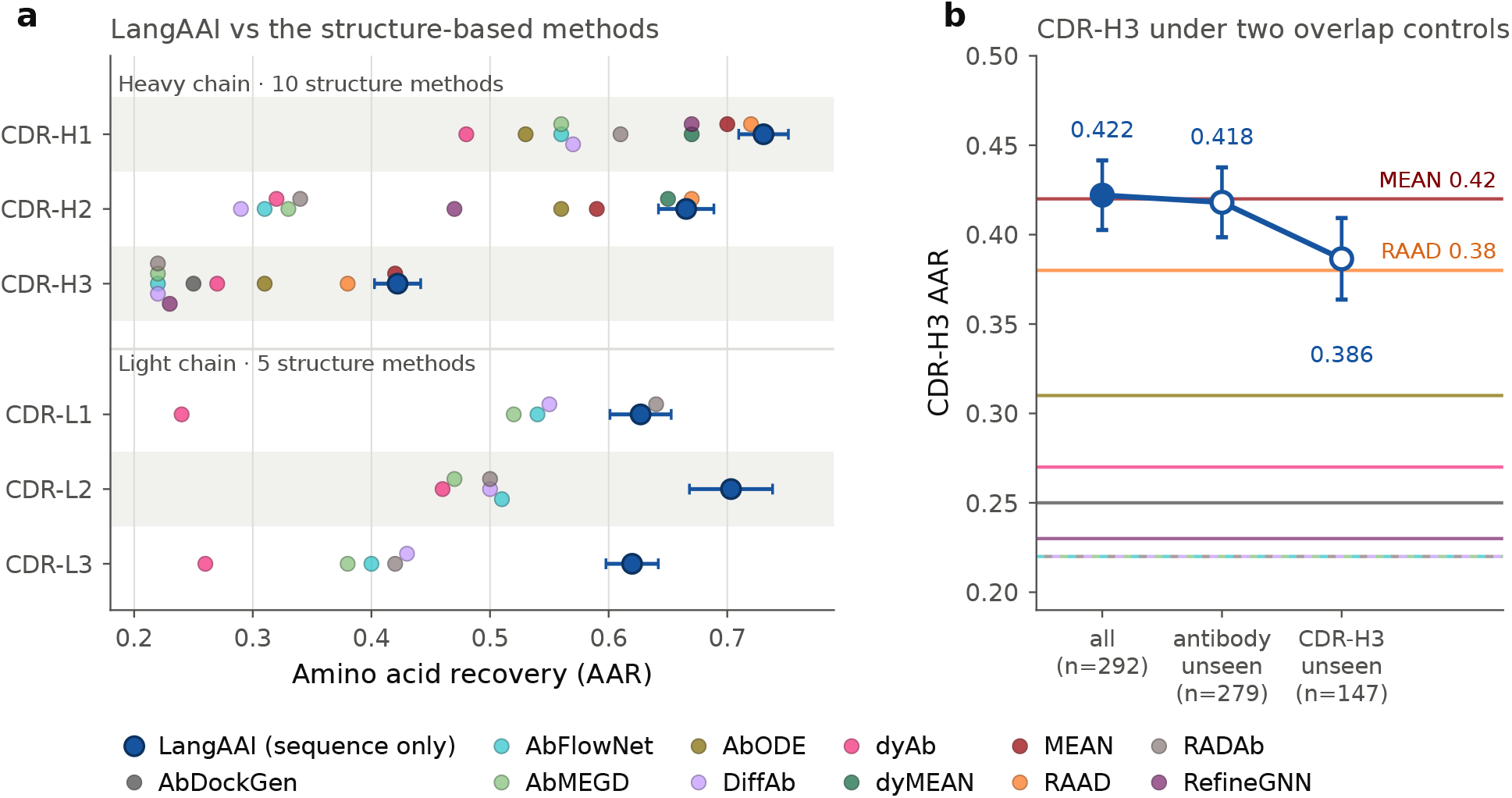
Comparison of LangAAI with structure-based CDR design on the CHIMERA-Bench’s own test split and metric. **(a)** Amino acid recovery (AAR) by CDR; each open marker is one structure-based method’s published mean, the blue marker is LangAAI’s mean with a 95% confidence interval over complexes; the best-performing structure-based method is labeled in each row. **(b)** Sensitivity of the CDR-H3 result from two different definitions of training-set overlap; grey lines mark each of the ten structure-based CDR-H3 methods; LangAAI is shown unrestricted (filled, *n* = 292) and under each control (open), with 95% confidence intervals.

To address potential data leakage from the shared SAbDab [Dunbar et al., 2014] database, we evaluated sequence overlap under two criteria. Figure 3b shows CDR-H3 Amino Acid Recovery (AAR) in two different definitions of training-set overlap. At the whole-chain level, 4.5% (*n* = 13/292) of test heavy chains appear verbatim in our training set. Excluding these altered regional AARs by at most, leaving five of six rankings unchanged and shifting CDR-H3 nominally from first to second (0.422 *→* 0.418 vs. MEAN 0.42). At the individual loop level, sequence redundancy is predictably high for conserved regions and so exact-match novelty is only biologically interpretable for the hypervariable CDR-H3 (49.7% overlap). Restricting analysis to the 147 complexes with strictly unseen CDR-H3 loops produced an AAR of 0.386 (95% CI 0.364–0.409). This secures second place among all eleven models, trailing RAAD (0.38) by less than 0.01. This slight decrement is independent of loop length, which is functionally identical across subsets (15.1 vs. 15.9 residues).

Taken all together, these results demonstrate that LangAAI can achieve competitive sequence-recovery capabilities of state-of-the-art, 3D-conditioned models using only sequence data. This comparative analysis evaluates informational sufficiency rather than establishing a definitive model hierarchy. While structural data remains a highly informative modality, high Amino Acid Recovery (AAR) does not physically guarantee that a designed loop will properly fold or functionally bind its intended target.

### 2.5. Generative sequence probabilities reflect experimental affinity maturation landscapes

LangAAI computes a probability distribution over the candidate amino acids at every masked antibody position. To determine whether these computational distributions reflect physical binding realities, we validated the model against independent, experimentally derived affinity data. We utilized two published AlphaSeq affinity-maturation datasets [Agarwal et al., 2025], which provide *in vitro* binding measurements for specific antibody mutations evaluated against a fixed target: (i) the pembrolizumab-scFv against PD-1 (YM_0852) and (ii) the VHH72 nanobody against the SARS-CoV-2 receptor-binding domain (YM_0549). By conditioning LangAAI strictly on the parent antibody and the target antigen, we quantified how accurately the model’s generated amino acid distributions align with these ground-truth experimental affinity landscapes.

For each dataset, we took the parent (“seed”) antibody together with its cognate antigen and identified the heavy-chain CDR positions that the library had mutagenized (24 for PD-1 and 34 for CoV2-RBD). We masked each position in turn, passed the masked antibody–antigen pair through LangAAI, and extracted a discrete probability distribution, i.e., *p*(*a* |masked position, remaining sequence, antigen), across the twenty canonical amino acids. Each experimentally measured variant was then assigned a score corresponding to the cumulative log-probability of its specific substitutions (Σlog *p*). In the case of a double mutant, this process composes two independently computed single-position conditionals and therefore neglects epistasis. We compared these scores against all library variants whose substitutions were entirely confined to the scored window (yielding 432 single and 4,460 double mutants for PD-1; 607 single and 1,870 double mutants for CoV2-RBD). Because affinity is reported as log_10_(*K*_*d*_), improved binding corresponds to a numeric decrease. We defined a variant as *improved* when its affinity score decreased more than two times the standard errors below the seed’s 50-replicate mean, i.e., beyond the measurement noise in the assay itself, not merely below the seed’s average by chance. Restricting the comparison population to the scored positions isolated residue choice from position choice.

Figure 4a and Figure 4b show that model scores and measured binding affinity were significantly associated in both datasets. For double mutants, we observed Spearman correlations of *ρ* = 0.243 (*P* = 6.9 *×* 10^*−*61^; *n* = 4, 460) for PD-1 and *ρ* = 0.232 (*P* = 2.9 *×* 10^*−*24^; *n* = 1, 870) for CoV2-RBD, with the running median of affinity gain steadily rising across the score range. This relationship was graded rather than threshold-like. When partitioned into quintiles based on log *p*, the proportion of variants outperforming the seed beyond the noise floor (i.e., decreased more than two times standard error below the replicate mean) increased consistently. For PD-1, this proportion rose at every quintile, growing from 1.7% to 13.5%, representing a shift from 0.21*×* to 1.69*×* the 7.9% base rate. For CoV2-RBD, the proportion climbed from 6.1% to 16.6% across the first four quintiles before leveling off at 16.0% in the top quintile, compared to a 12.1% base rate (Figure 4c). Correspondingly, mean affinity improved monotonically across these bins in both datasets, by 0.79 and 0.43 log_10_(*K*_*d*_) units, respectively (Figure 4d). Finally, ranking variants by probability alone separated improved variants from the remainder with an AUC of 0.665 for PD-1 and 0.616 for CoV2-RBD (Figure 4e).

**Figure 4:**
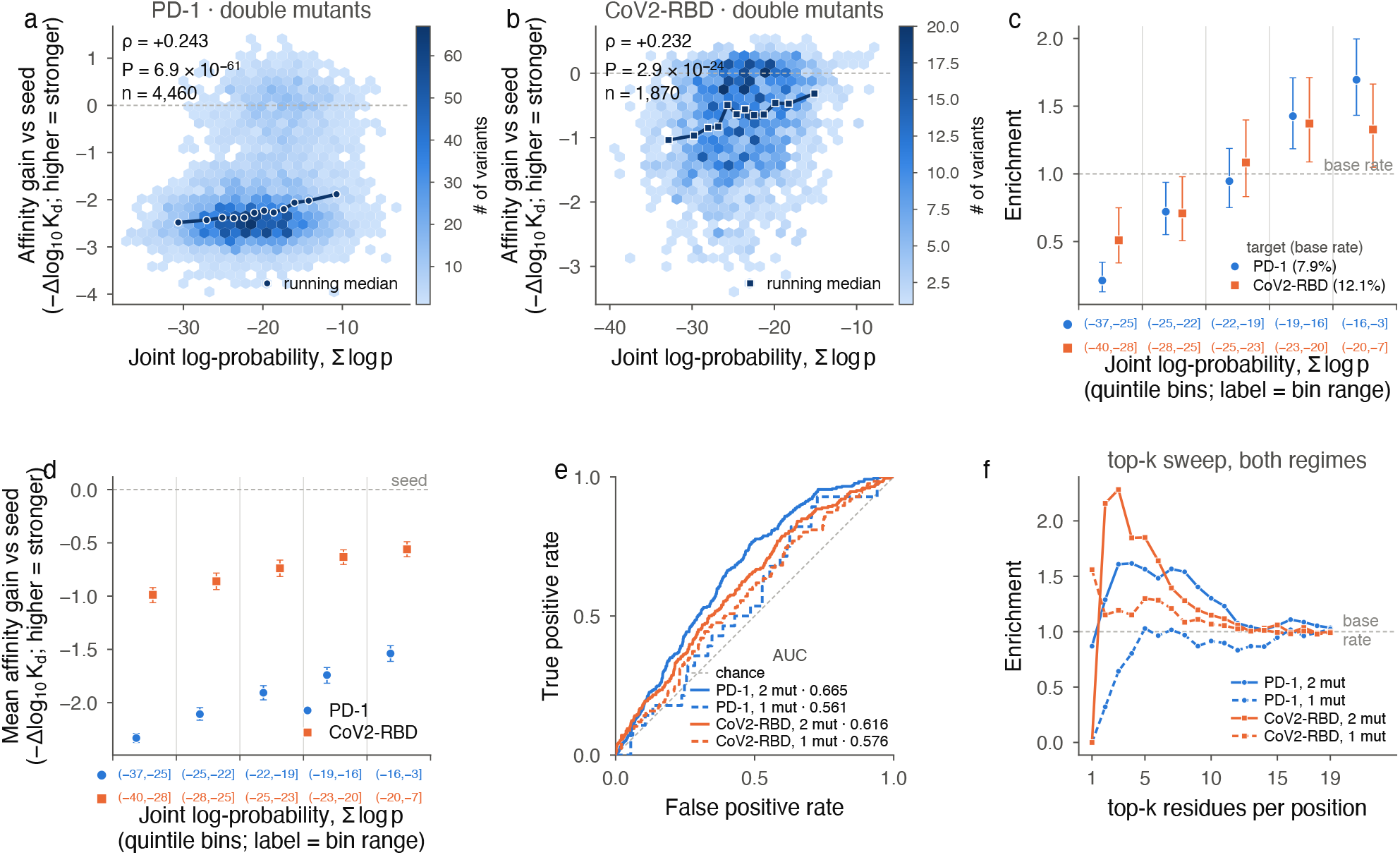
LangAAI provided masked-residue probability ranks antibody variants by experimentally measured affinity. **(a, b)** Joint log-probability against measured affinity gain for PD-1 (**a**, *n* = 4,460 variants) and CoV2-RBD (**b**, *n* = 1,870); hexagonal bins are variant counts, on independent color scales between the two panels; the dark line is the running median over 12 equal-count score bins; *ρ* and P-values are two-sided Spearman rank correlation. **(c)** Fraction of variants whose affinity is more than two times the standard errors below the seed’s 50-replicate mean, expressed as enrichment over each dataset’s own base rate; error bars are 95% Wilson score intervals. **(d)** Mean affinity gain per quintile for the same bins; error bars, 95% percentile bootstrap intervals. **(e)** Receiver operating characteristic for identifying improved variants by joint log-probability; solid are double mutants; dashed are single mutants; AUC computed from the Mann–Whitney U statistic. **(f)** Enrichment over the base rate among variants reachable by proposing the top-k highest-probability residues at every position independently.

We observed a similar but weaker association in the single-mutant regime (*ρ* = 0.214 and 0.216, *P* = 7.5 *×* 10^*−*6^ and 7.8 *×* 10^*−*8^; AUC 0.561 and 0.576; see Supplementary Figure 2a and Supplementary Figure 2b). This discrepancy is driven by two structural limitations rather than a difference in true model performance: single substitutions inherently span a narrower score range than double mutations (10.9 versus 15.3 log units across quintile means for PD-1), and the dataset contains an order of magnitude fewer single mutants.

Figure 4f shows that applying a per-position proposal budget successfully concentrated improved variants at shallow sampling depths. To evaluate this, we ranked all non-wildtype residues per position, retaining a variant at depth *k* only if every constituent substitution ranked within the top *k* for its respective site. We then compared the success rate of these selected variants against the baseline success rate of the unconstrained library (PD-1: 6.5% for singles, 7.9% for doubles; CoV2-RBD: 20.8% for singles, 12.1% for doubles). Across all 76 combinations of depth, dataset, and mutation regime, the success rate of the constrained variants never fell significantly below the library base rate (minimum lower-tail *P* = 0.15, hypergeometric test). This confirms that ranking by probability consistently matches or outperforms random library sampling. Enrichment above the baseline peaked strongly at *k* = 3 *to* 6. For double mutants, PD-1 enrichment reached 1.62*×* the base rate at *k* = 4 (with 12.8% of 304 constrained variants acting as reliable improvers), while CoV2-RBD peaked at 2.28*×* at *k* = 3 (27.6% of 58 variants). Both datasets regressed to their baseline rates by *k* ≈ 12, indicating that a restrictive budget of a few residues per position captures a concentrated signal that broader inclusion dilutes. Finally, because baseline success rates inherently vary, relative enrichment and absolute yield emphasize different regimes. CoV2-RBD single mutants exhibited only a 1.56*×* relative enrichment at *k* = 1, yet produced the highest absolute hit rate of any dataset, yielding 32.4% from 34 retained variants.

Collectively, these results demonstrate that the masked-residue probabilities generated by LangAAI effectively rank antibody variants according to their empirical binding affinity. Prioritizing a candidate pool using joint log-probability strongly concentrates improved variants within the highest-scoring fractions. Specifically, the proportion of improved variants increases approximately 8-fold from the bottom to the top quintile for PD-1 (1.7% to 13.5%), and 2.6-fold for CoV2-RBD (6.1% to 16.0%). Overall, this probability-based ranking successfully discriminates affinity-enhancing variants from the background population, yielding an AUC of 0.665 and 0.616, respectively.

### 2.6. LangAAI’s residue likelihood generalises to an independent binding-affinity benchmark

To test whether LangAAI’s antigen-conditioned likelihood tracks binding affinity rather than sequence plausibility alone, we evaluated it on the AbBiBench [Zhao et al., 2025] dataset. AbBiBench is a benchmark of 16 antibody–antigen systems, and it features variants of a parent antibody with experimentally measured affinities. In the AbBiBench study, the authors evaluates 19 methods including ESM-2 [Lin et al., 2023], AntiBERTy [Ruffolo et al., 2021], SaProt [Su et al., 2023], ProSST [Li et al., 2024], ESM-3 [Hayes et al., 2025], CurrAb [Burbach and Briney, 2025], ProGen2 [Nijkamp et al., 2023], ProtGPT2 [Ferruz et al., 2022], ProteinMPNN [Dauparas et al., 2022], ESM-IF [Hsu et al., 2022], AntiFold [Høie et al., 2025], DiffAb [Luo et al., 2022], MEAN [Kong et al., 2022], dyMEAN [Kong et al., 2023] and two biophysics based models. We scored every variant under the benchmark’s own likelihood definition, *logP*(*s*) =Σ_*i*_ *logP*(*s*_*i*_ *s, x*) where *s* is the antibody and *x* is the antigen, a single forward pass over the uncorrupted complex. These scores were then correlated with empirical binding affinities using Spearman’s rank correlation (*ρ*). This represents a strict zero-shot evaluation as no LangAAI parameters were tuned on these experimental data, and no benchmark variant is present within our training corpus.

Figure 5a shows that across the 16 systems LangAAI reached a mean Spearman *ρ* of 0.188, ranking fourth among all the methods and placing above every sequence-only model on the benchmark. The three methods that had a better mean Spearman, namely ProteinMPNN (0.299), ESM-IF1 (0.282) and AntiFold (0.207), are inverse-folding models that use the backbone coordinates. LangAAI exceeded ESM-2 by Δ*ρ* = 0.170 (Wilcoxon signed-rank *W* = 26, *P* = 0.029, 16 paired systems) and AntiBERTy by Δ*ρ* = 0.295 (*W* = 28, *P* = 0.039), with a smaller margin over the next sequence-only method, CurrAb (Δ*ρ* = 0.113, *W* = 35, *P* = 0.093). Moreover, despite lacking 3D coordinate data, LangAAI achieved statistical parity with the inverse-folding AntiFold model (Δ*ρ* = *−*0.019; *W* = 56, *P* = 0.56), remaining definitively behind ProteinMPNN (Δ*ρ* = *−*0.111; *W* = 17, *P* = 0.0063).

**Figure 5:**
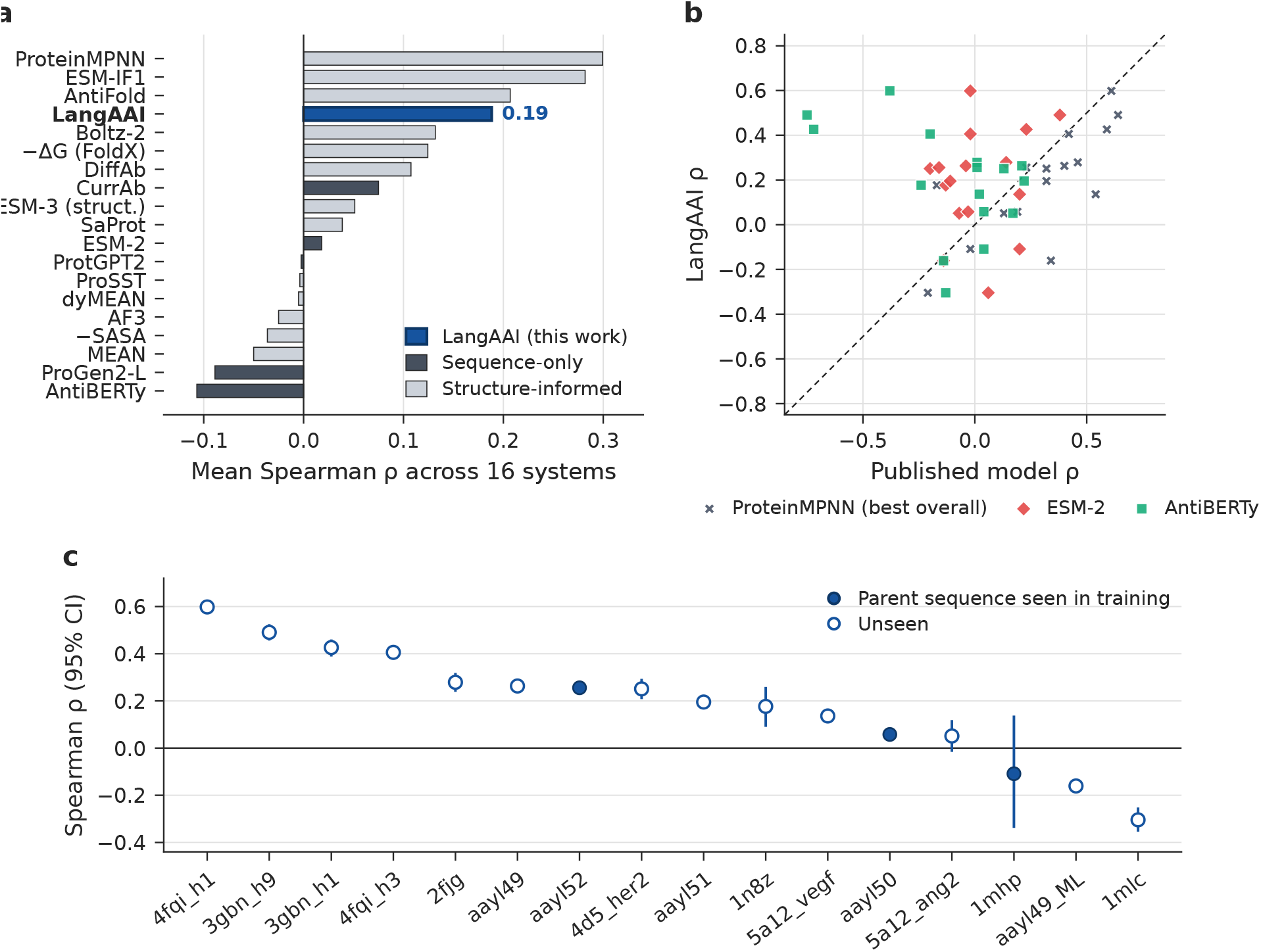
LangAAI on the AbBiBench binding-affinity benchmark. **(a)** Mean Spearman correlation between model log-likelihood and measured binding affinity across the 16 AbBiBench systems, for LangAAI and the 18 methods the benchmark reports. **(b)** Per-system correlation for LangAAI against three reference methods; each point is one system and the dashed line is the identity. Points above the line are systems on which LangAAI ranks variants better. **(c)** LangAAI’s correlation per system, ordered, with percentile bootstrap 95% confidence intervals over variants. Filled markers denote systems whose parent heavy chain was seen in LangAAI’s training split.

**Figure 6:**
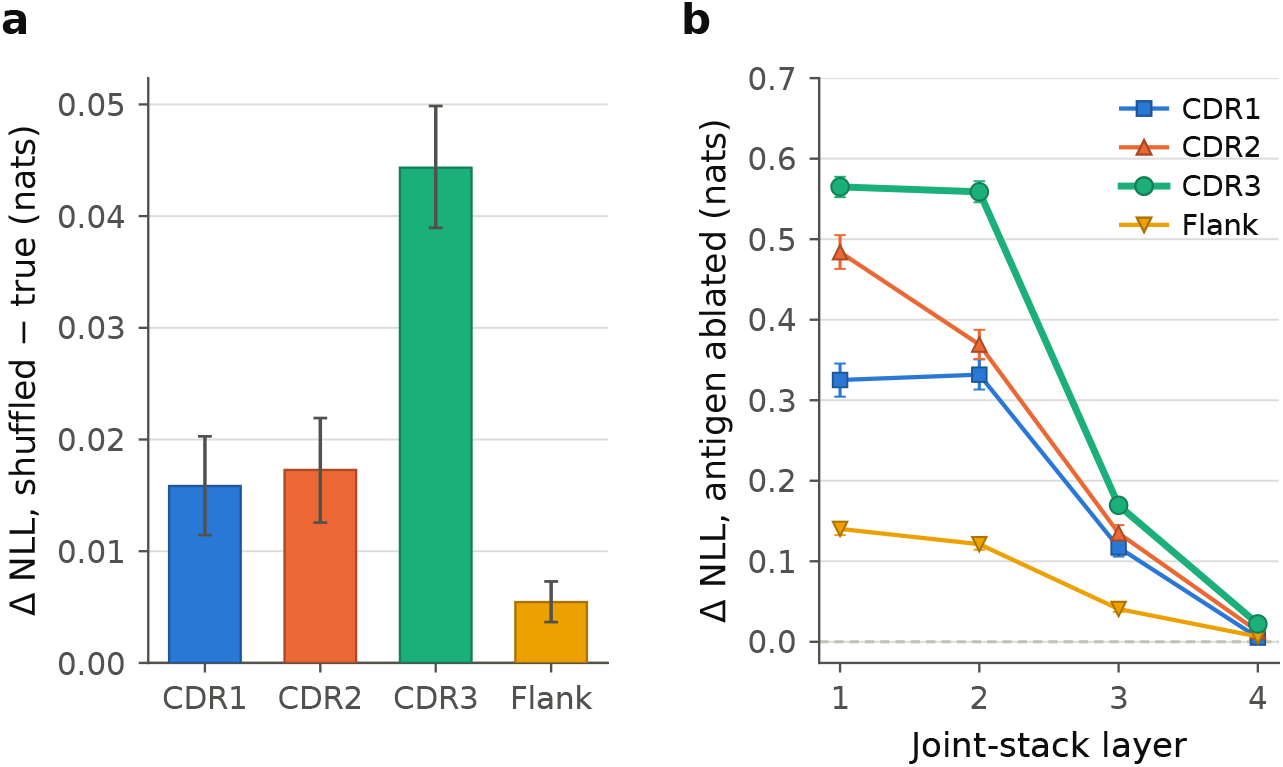
Antigen dependence is concentrated in CDR3. **(a)** Δ*NLL* between shuffled and true antigen per antibody region.**(b)** Δ*NLL* when the antigen’s contribution is ablated from one joint-stack layer onward.

Figure 5b and Figure 5c show that per-system performance exhibited strong heterogeneity. LangAAI performed best on the influenza haemagglutinin systems, achieving *ρ* = 0.598 (95% CI:0.593–0.603, *n* = 65, 094) for `4fqi_h1` and *ρ* = 0.491 (95% CI:0.456–0.525, *n* = 1, 842) for `3gbn_h9`. However, the correlation inverted significantly on two test complexes. For `1mlc` (*ρ* = *−*0.304, 95% CI:*−*0.354 to *−*0.252, *n* = 1, 229), this structural failure was shared by explicit backbone baselines ESM-IF1 ( −0.31) and AntiFold ( −0.31). In contrast, for `aayl49_ML` (*ρ* = *−*0.161, 95% CI: *−*0.182 to *−*0.139, *n* = 8, 953), ProteinMPNN maintained a positive correlation (*ρ* = 0.34). The benchmark contained two further non-informative systems: `5a12_ang2` yielded a statistically null correlation (*ρ* = 0.051, 95% CI: *−*0.016 to 0.119, *n* = 944), and `1mhp` (*ρ* = *−*0.109, 95% CI: *−*0.338 to 0.138) lacked sufficient statistical power (*n* = 69) to draw definitive conclusions regarding model efficacy.

Finally, because several benchmark antibodies are well-characterized therapeutics, we investigated whether this performance could stem from memorization. The evaluation is “zero-shot” in both data and objective: all 213,685 scored variants are absent from our training split by construction, no parameter was fitted to these data, and the task of ranking variants by affinity differs from the masked-reconstruction objective LangAAI was trained on. Besides, antigen provenance yielded no material impact on performance. Results are shown in Figure 5c. Systems with antigens absent from the training set averaged *ρ* = 0.177 (*n* = 8 systems), comparable to *ρ* = 0.196 for those with present antigens (*n* = 9 systems). Notably, the highest-performing family, `3gbn` (*ρ* = 0.426 and 0.491), involves an antigen strictly novel to training data. Performance was also not concentrated on the three complexes featuring a verbatim parent heavy chain in the training split (mean *ρ* = 0.068; Figure 5c, filled markers). Therefore, LangAAI’s predictive capability reflects true out-of-distribution generalization.

### 2.7. Antigen dependence localizes to CDR3

To characterize how the antigen influences reconstruction, we evaluated all 17,983 cognate test pairs, decomposing each by antibody region and by joint-stack layer. The majority of these data (69%) derive from patents, with the remainder sourced from HIV, `SARS-CoV-2`, literature, GenBank, and structural databases. Regions are labeled without distinguishing heavy from light chain, so the antigen dependency scores were pooled across CDR1, CDR2 and CDR3 when a light chain is present. However, as 84.7% of the evaluated antibodies are heavy-chain only, these regions overwhelmingly reflect heavy-chain sequences. To decompose the antigen’s mechanistic influence across specific antibody regions and joint-stack layers, we applied two distinct tests: (i) a shuffled-antigen perturbation evaluated independently for each target region, and (ii) a layer-local causal ablation that zeroes the antigen’s hidden state prior to its entry into each of the four joint-stack layers.

Substituting a non-cognate antigen degraded CDR3 reconstruction the most (ΔNLL = 0.044 nats, 95% CI [0.039 *−* 0.050]). This degradation was 2.6-fold greater than in CDR2 (0.017, [0.013, 0.022]) and 2.8-fold greater than in CDR1 (0.016, [0.011, 0.020]). The framework flank exhibited the smallest response, remaining measurable but minimal (0.005, [0.004, 0.007]). This flank response was eight-fold lower than that of CDR3, corresponding to a perplexity ratio of 1.005 compared to 1.045 for CDR3. Such stability is expected, given that flanking residues are largely predictable from the local antibody sequence alone. Furthermore, CDR3 was the primary region where the attention mechanism itself responded to the substitution. The shift in attention mass correlated with the change in reconstruction loss significantly more strongly for CDR3 (Pearson *r* = 0.28 to 0.29 across all four layers) than for CDR1 (0.09 to 0.11), the framework flank (0.07 to 0.11), or CDR2 (0.03 to 0.04) (more details in Appendix, Section G). Outside of CDR3, these correlations remain weak and do not track the loss hierarchy. Consequently, this attention analysis reinforces the unique target sensitivity of CDR3 rather than a broad region-wise correspondence.

Removing the antigen entirely rather than substituting it incurred a substantially higher cost. For CDR3, this ablation penalty reached 0.565 nats (95% CI [0.552, 0.577], *n* = 17,760) at the first layer. This was the largest effect observed for any region at any layer, and it fell to 0.022 nats ([0.021, 0.024]) by the fourth layer. Because the antigen is concatenated into the joint sequence once before the first layer, ablating layer *i* also removes its content from every subsequent layer. Consequently, this decay largely reflects how much of the forward pass each ablation eliminates. Crucially, the ablation reproduced the region-specific ordering of the antigen swap. CDR3 exhibited the largest effect at every layer, while the framework flank showed the smallest at 0.140 nats ([0.133, 0.148]), which is four-fold below CDR3. This confirms the effect is region-specific rather than a generic consequence of dropping an input stream.

## 3. Discussion

Our results demonstrate that antigen-conditioned modeling of antibody hyper-variable regions does not require explicit 3D coordinates. Instead, it demands a training objective that computationally forces the antigen to dictate the masked residue. Under that objective, a lightweight joint transformer over frozen sequence encoders recovers CDR residues more accurately than antibody-specific language models. Furthermore, it measurably changes its predictions when the cognate antigen is replaced, matches structure-conditioned design methods on their own recovery benchmark, and ranks experimentally measured variants by affinity well on an external benchmark. None of achievements rely upon an input backbone, a docking pose, or a predefined epitope.

A broader implication of our work is that simply providing a model with the antigen sequence does not guarantee genuine antigen conditioning. Although MINT and PLM-interact both read the target sequence, their generative outputs were statistically indistinguishable from an antigen-blind baseline across all tested partitions. This reflects their underlying training objectives, which optimize for discriminating interacting pairs rather than reconstructing hyper-variable binding loops. The critical consequence is that baseline reconstruction accuracy cannot prove a model actually utilizes its conditioning variable. Because antibody frameworks and local sequence statistics are inherently predictable, a model can achieve high recovery metrics while entirely bypassing the target antigen. The paired-shuffle difference introduced here provides a computationally efficient solution that requires no retraining and strictly returns zero for any model ignoring its conditioning input. We propose that any architecture claiming conditional generation on paired sequences report a comparable contrast metric alongside standard accuracy.

The structural localization of this antigen dependence provides crucial insights beyond its raw magnitude. Two distinct perturbations, namely replacing the antigen with a non-cognate decoy and zeroing its hidden state prior to a given joint layer, independently localize the effect to CDR3. Conversely, these interventions leave the framework flank nearly untouched (0.034 versus 0.001 nats under substitution). This pattern strictly aligns with the biophysical mechanics of target recognition rather than baseline sequence statistics, because framework positions are largely recoverable from local germline context while the hyper-variable CDR3 is not. The convergence of a distributional perturbation and a causal ablation provides stronger evidence than either test alone. Together, they demonstrate that the antigen pathway in this architecture performs localized, structurally specific work rather than merely supplying a diffuse background signal.

In practice, LangAAI’s capability facilitates rapid candidate prioritization. Given a parent antibody and a target sequence, our method ranks proposed substitutions prior to the commitment of experimental assay capacity. On the AbBiBench benchmark, the model achieves statistical parity with AntiFold, a coordinate-reading inverse-folding architecture (Δ*ρ* = *−*0.019, *P* = 0.56), although it remains behind ProteinMPNN. Furthermore, across two AlphaSeq maturation campaigns, restricting evaluation to the top-ranked fraction concentrates improved variants by 1.6-to 2.5-fold relative to the unfiltered library base rate. These margins provide a highly effective triage step. The primary operational advantage is that these predictions are accessible entirely without a solved structural complex. Because fewer than 8,000 antibody–antigen structures exist and predictive docking algorithms yield accurate interfaces in roughly one out of ten cases, structure-conditioned ranking is frequently unreliable or entirely unavailable for novel therapeutic targets. We note explicitly that this evaluates a predictive ranking capability, as no sequences generated by the model were synthesized in this study.

## 4. Materials and Methods

### 4.1. Data

We used antibody–antigen pairs curated from patent filings, structural depositions, and published binding assays from the ASD database [Czerwiński et al., 2026]. We retained only positively-labeled pairs with a heavy-chain sequence present, yielding 172,537 pairs. To prevent identity-driven memorization from being mistaken for antigen-aware generalization, we independently clustered antibody and antigen sequences using MMseqs2 [Steinegger and Söding, 2017] (90% identity threshold) and split into train/validation/test by cluster membership, yielding 139,644/14,617/18,276 pairs accordingly (full breakdown in Table 1). We further stratified the validation and test splits by cluster novelty relative to training, yielding three regimes of increasing difficulty: (i) `ab_novel`, in which the antibody cluster is absent from training but the antigen may have been seen; (ii) `ag_novel`, in which the antigen cluster is absent from training but the antibody may have been seen; and (iii) `both_novel`, in which neither cluster appears in training. The antibody-held-out category (`ab_novel`) was the primary evaluation set, since an unseen antibody cluster carries no identity signal the model could have memorized, ensuring that performance does not depend on antibody identity.

**Table 1:** Composition of the entity-holdout data partition.

| Split | Category | Pairs | Unique antibodies | Unique antigens |
| --- | --- | --- | --- | --- |
| Train | — | 139,644 | 31,675 | 7,410 |
| Validation | ab_novel | 6,950 | 1,302 | 2,220 |
| Validation | ag_novel | 7,348 | 4,432 | 335 |
| Validation | both_novel | 319 | 146 | 107 |
| Test | ab_novel | 8,551 | 1,887 | 2,030 |
| Test | ag_novel | 8,804 | 5,154 | 474 |
| Test | both_novel | 628 | 277 | 170 |

### 4.2. Model Architecture

LangAAI (i) encodes the antibody and antigen with two independent frozen protein language models, (ii) projects both into a shared representation space, (iii) concatenates them behind a learned classification token, and (iv) processes the joint sequence with an unrestricted self-attention stack. A masked-residue prediction head, tied to the antibody encoder’s own vocabulary embedding, was evaluated on the antibody segment of the output only. An overview of LangAAI’s architecture is shown in Figure 7.

**Figure 7:**
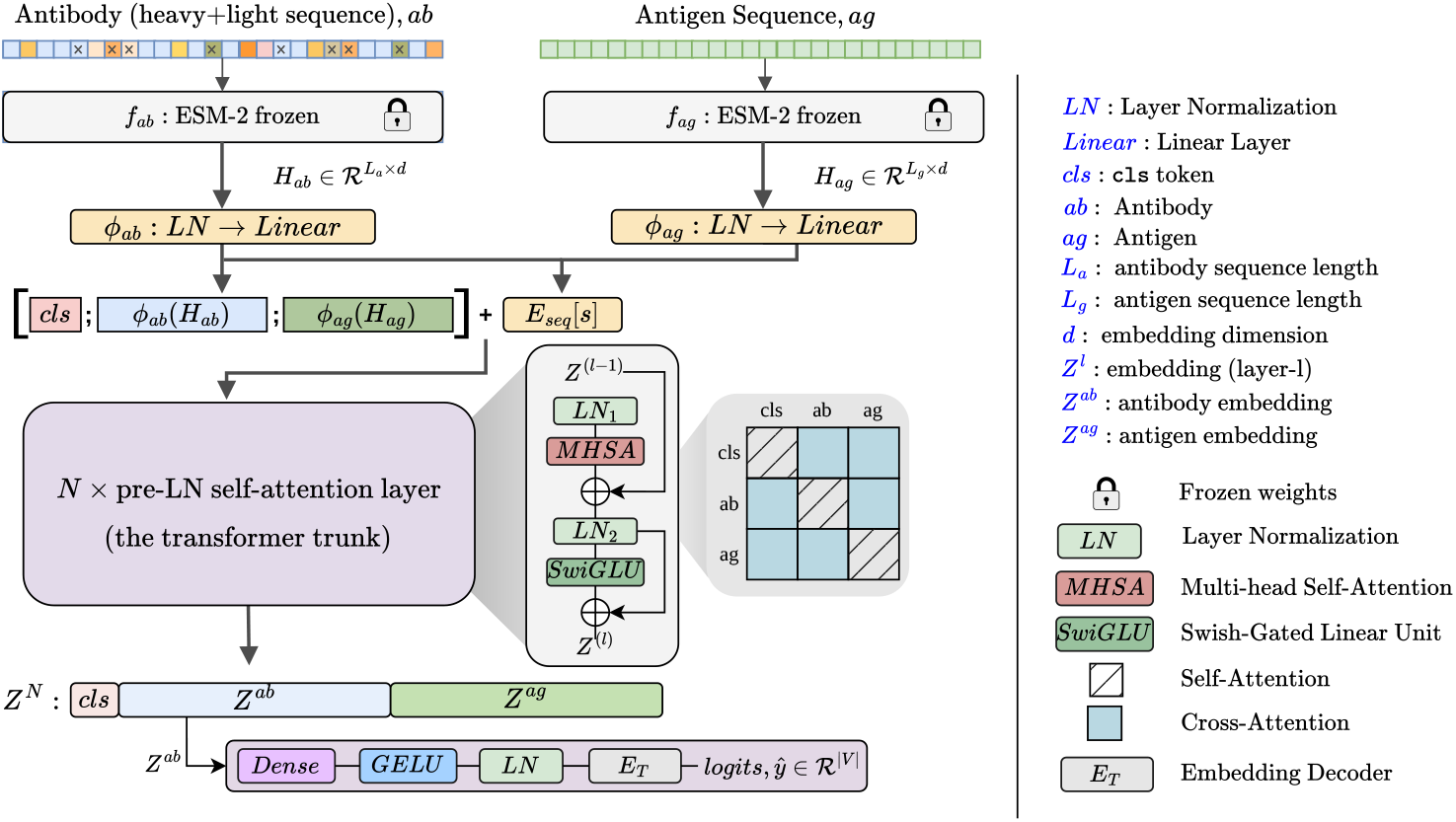
LangAAI architecture. Two frozen ESM-2 [Lin et al., 2023] encoders process the (masked) antibody and antigen sequences independently; trainable adapters project both into a shared space; a classification token and segment embeddings assemble the joint sequence, processed by *N*=4 layers of unrestricted self-attention; a tied prediction head, sharing weights with the frozen antibody encoder’s embedding decoder, is evaluated on the antibody segment only.

#### Sequence encoding

In LangAAI, the antibody sequence **a** (heavy chain, concatenated with the light chain when present) and the antigen sequence **g** are each encoded with an ESM-2 protein language model [Lin et al., 2023], which is kept frozen throughout training.

We obtain the embeddings the antibody and the antigen, as follows

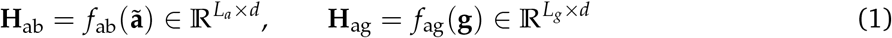

where **ã** is the masked antibody sequence and *d*=480 (ESM-2, 12 layers, 35M parameters). We used a smaller ESM-2 variant rather than one of the larger checkpoints, for two main reasons. First, the role of *f*_ab_ and *f*_ag_ in our architecture is to supply position-aware, evolutionarily informed context for each residue, not to resolve the binding signal itself, which is instead learned by the stack of joint co-attention layers. Second, the resulting embedding width *d*=480 keeps the cost of co-attention (*O*(*L*_*a*_*L*_*g*_*d*)) tractable for full-length antigens, permitting larger batches and avoiding the sequence truncation that a wider representation would force under the same memory budget. In other words, representational capacity is deferred to the components that are actually optimized on the antibody– antigen interaction objective, and ESM-2 is used strictly as a fixed contextual featurizer.

Each encoder’s output is then independently normalized and linearly projected.

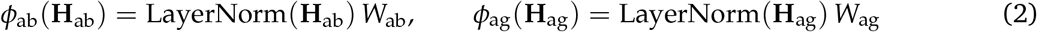

with *W*_ab_, *W*_ag_ ∈ ℝ^*d*×*d*^ each initialized to the identity matrix, so that training began from normalized, unrotated encoder features.

#### Joint sequence assembly

A learned classification vector **c** is prepended and a four-way segment embedding (0=classification token, 1=heavy chain, 2=light chain, 3=antigen) is added. The joint representation **Z**^(0)^ is defined as

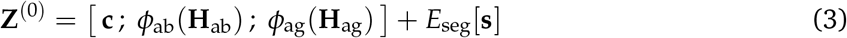

No positional encoding is added to the joint sequence because both encoders are already position-aware internally, and an absolute joint index would make the antigen’s encoded position depend on the antibody’s length, a relationship with no physical meaning.

#### Co-attention

The joint representation **Z**^(0)^ is processed by *N* pre-norm transformer layers with multi-head self-attention and a gated feed-forward sublayer.

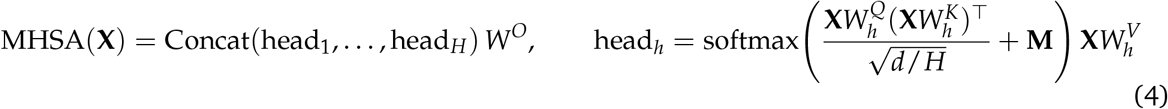

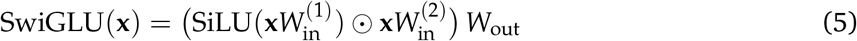

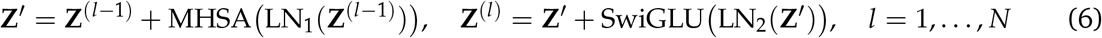

with *N*=4 layers, *H*=8 heads, feed-forward expansion ratio 4, and dropout 0.1. Critically, the additive mask **M** in Equation 4 encodes only sequence padding (*M*_*ij*_ = *−*∞ if and only if key position *j* was a padding token, for every query *i*) and impose no other structure. Every antibody position could attend to every antigen position and vice versa, under one shared set of attention weights, with no separate parameters for cross-chain interactions. This unrestricted formulation is the central architectural choice under evaluation.

#### Masked-residue prediction head

The output projection is tied to *E*, the antibody encoder’s own token embedding matrix (i.e., the identical parameter tensor, held frozen), and evaluated only over antibody positions *i ∈ {*1, …, *L*_*a*_*}* of the joint output.

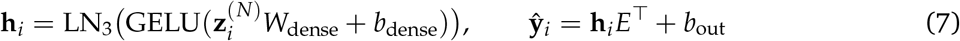

We reuse the pretrained embedding matrix *E* in the output layer, allowing the model to inherit ESM-2’s rich amino acid geometry. At initialization, this ensures the head already recognizes which residues are substitutable at any given position. This gives the randomly initialized joint stack a significant head start. Rather than having to relearn basic amino acid biochemistry from scratch, it only needs to learn how the antigen’s presence shifts this pre-existing baseline distribution.

### 4.3. Training

#### Masking scheme

To construct the training task, we masked antibody positions independently based on their structural region. We targeted 40% of the residues in CDR3, 15% in CDR1, CDR2, and the flanks (near-CDR region). Then the masked positions were corrupted following the standard 80/10/10 masked-LM recipe [Devlin et al., 2019]. The higher masking rate for CDR3 was a critical design choice. If we had used a standard 15% rate, the model could rely on the remaining 85% of the loop to fill in the blanks, effectively bypassing the antigen altogether. Our 40% rate forced the model to utilize the antigen context, resulting in about 12 masked tokens per sequence. Finally, to fit our maximum sequence length budget (*L*^max^), we dynamically cropped the antigen to a window of width min(*L*_*g*_, max(1, *L*^max^ *−L*_*a*_ *−*1)), applying a random crop offset during training and a centered crop during evaluation.

#### Loss function and Optimization

The training minimized cross-entropy over masked positions only, thus the loss function was defined as

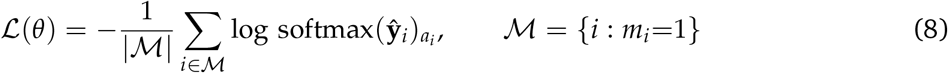

We optimized the model parameters using AdamW with a learning rate of 3 *×* 10^*−*4^ and a weight decay of 0.01. To prevent over-regularization, we excluded all LayerNorm parameters, biases, the classification token, and the segment embedding from weight decay. The learning rate itself was governed by a cosine decay schedule, preceded by a linear warm-up phase during the first 5% of training steps.

#### Model selection and evaluation metric

Relying on absolute masked-residue loss is inherently flawed due to sequence memorization. In fact, 44% of our primary test set (`ab_novel`) features CDR3 sequences seen verbatim during training. To eliminate this bias and isolate the antigen’s actual contribution, we evaluated each test example against a strictly paired counterpart. By holding the antibody and masked positions constant while substituting a shuffled antigen, we compared the model’s likelihood across both scenarios.

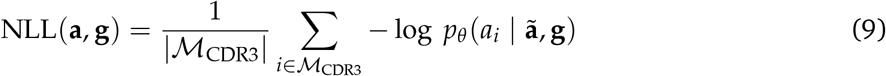

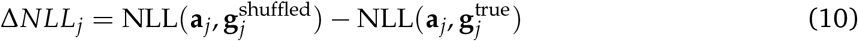

Any performance boost from memorized antibody structure affected both terms in Equation 10 and perfectly canceled out. As a result, Δ*NLL*_*j*_ captured solely the antigen’s marginal contribution, ensuring that an antigen-blind model yielded exactly zero by mathematical design rather than by empirical observation. We generated the shuffled pairs by drawing mismatched antigens from the same evaluation category, excluding any sequences that shared a cluster with the true antigen.

We also measure top-1 accuracy, or Amino Acid Recovery (AAR). This tracks the precise fraction of masked residues successfully restored via arg max decoding. To ensure balanced representation, we calculated this recovery rate on a per-example basis before taking the global average.

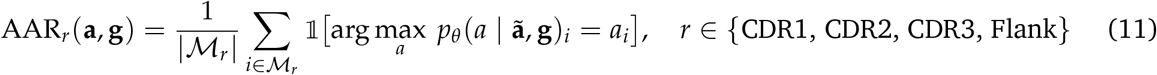

