## Supplementary File for "Sequence-only antigen context improves modeling of antibody hypervariable regions"

### Supplementary Material

**Faisal Bin Ashraf, Stefano Lonardi**

Department of Computer Science and Engineering, University of California, Riverside

**Source Code:** https://github.com/fbabd/LangAAI

#### 1. Notation

Symbols as introduced, in order of first appearance in Materials and Methods.

Table 1: Mathematical symbols and definitions.

| Symbol | Definition |
|---|---|
| $\mathbf{a} = \mathbf{a}_H \oplus \mathbf{a}_L$ | Antibody sequence; heavy chain concatenated with light chain when present |
| $\mathbf{g}$ | Antigen sequence |
| $L_a, L_g$ | Antibody and antigen sequence lengths (post-crop for $L_g$) |
| $r_i$ | Region label of antibody position $i \in \{\texttt{CDR1}, \texttt{CDR2}, \texttt{CDR3}, \texttt{FLANK}, \texttt{NONE}\}$ |
| $m_i$ | Mask indicator at position $i$ |
| $\tilde{\mathbf{a}}$ | Corrupted (masked) antibody sequence |
| $P_{\text{ab}}, P_{\text{ag}}$ | Antibody-/antigen-cluster pool assignment (train/validation/test) |
| $f_{\text{ab}}, f_{\text{ag}}$ | Antibody and antigen ESM-2 encoders |
| $d$ | Model width, pinned to the frozen encoder's hidden size ($d$=480) |
| $\mathbf{H}_{\text{ab}}, \mathbf{H}_{\text{ag}}$ | Encoder outputs |
| $\phi_{\text{ab}}, \phi_{\text{ag}}$ | Per-side cross-modal projections (Eq. 2) |
| $W_{\text{ab}}, W_{\text{ag}}$ | Projection weight matrices, identity-initialized |
| $\mathbf{c}$ | Learned classification-token vector |
| $E_{\text{seg}}, \mathbf{s}$ | Segment embedding table and per-position segment id |
| $\mathbf{Z}^{(l)}$ | Joint sequence representation after self-attention layer $l$ |
| $N, H$ | Self-attention depth and head count ($N$=4, $H$=8) |
| $\mathbf{M}$ | Additive attention mask (padding only; Eq. 4) |
| $E$ | Frozen antibody-encoder token embedding matrix, reused by the tied head |
| $\hat{\mathbf{y}}_i$ | Predicted logits at antibody position $i$ |
| $\rho(r_i)$ | Region-dependent masking rate (Eq. 10) |
| $L^{\text{max}}$ | Joint sequence length budget |
| $\theta$ | All trainable parameters |
| $\mathcal{M}, \mathcal{M}_{\text{CDR3}}$ | Set of masked positions; subset restricted to CDR3 |
| $\text{NLL}(\mathbf{a}, \mathbf{g})$ | Mean negative log-likelihood over $\mathcal{M}_{\text{CDR3}}$ (Eq. 14) |
| $\Delta_j$ | Paired shuffled-minus-true NLL difference for example $j$ (Eq. 15) |

#### 2. Source composition of the modeling dataset

Composition of the 172,537 antibody–antigen pairs used after entity-holdout routing (Materials and Methods, Data; Table 1 of the main text), by originating data source.

Table 2: Distribution of data sources for paired sequences.

| Source | Pairs | % of total | Cumulative % |
|---|---|---|---|
| Patent filings | 136,524 | 79.13 | 79.13 |
| COVID-19 structural/serological studies | 13,074 | 7.58 | 86.70 |
| HIV structural/serological studies | 11,957 | 6.93 | 93.63 |
| Published literature | 5,410 | 3.14 | 96.77 |
| GenBank | 1,928 | 1.12 | 97.89 |
| MET-antigen collection | 1,515 | 0.88 | 98.77 |
| Structural depositions (antibodies) | 1,503 | 0.87 | 99.64 |
| Structural depositions (nanobodies) | 592 | 0.34 | 99.98 |
| Other (SKEMPI v2, AbDesign, AB-Bind) | 34 | 0.02 | 100.00 |

Patent-derived sequences dominate the corpus (79.1%). This is discussed as a limitation on data diversity in Supplementary Note 2, since patent antibody sequences are disproportionately drawn from combinatorial-library variants of a small number of parent scaffolds.

#### 3. Model

The complete model contained 15,476,193 trainable parameters (Table 3), held atop a single frozen antibody encoder (33,269,521 parameters). A second, architecturally identical ESM-2 instance was used only to precompute the antigen cache (Data) and was not retained as part of the trained model or its forward graph.

Table 3: Parameter budget ($N$=4, $d$=480, $H$=8).

| Component | Parameters |
|---|---|
| Self-attention stack (4 layers + output norm) | 14,779,200 |
| Prediction head (excluding the tied embedding) | 231,873 |
| Cross-modal projections (both sides) | 462,720 |
| Segment embedding | 1,920 |
| Classification token | 480 |
| **Total trainable** | **15,476,193** |
| Frozen antibody encoder (within the model) | 33,269,521 |
| **Total, model object** | **48,745,714** |

#### 4. Masking scheme

For each antibody position $i \in \{1, \ldots, L_a\}$ with region label $r_i$, a mask indicator was drawn independently, as follows

$$m_i \sim \text{Bernoulli}\big(\rho(r_i)\big), \qquad \rho(r_i) = \begin{cases} 0.40 & r_i = \texttt{CDR3} \\ 0.15 & r_i \in \{\texttt{CDR1}, \texttt{CDR2}, \texttt{FLANK}\} \\ 0 & r_i = \texttt{NONE}, \end{cases} \tag{S1}$$

where FLANK denotes the residues immediately adjacent to each CDR and NONE the remaining framework positions. Selected positions were then corrupted following the standard BERT protocol [?],

$$\tilde{a}_i = \begin{cases} \texttt{[MASK]} & \text{with probability } 0.8 \\ u \sim \text{Uniform}(\text{amino acids}) & \text{with probability } 0.1 \\ a_i & \text{with probability } 0.1 \end{cases} \quad \text{if } m_i{=}1, \tag{S2}$$

and $\tilde{a}_i = a_i$ otherwise. Note that the loss in Equation (**??**) is evaluated over all positions with $m_i{=}1$, including the 10% left uncorrupted, so that the model cannot infer which positions are scored from the presence of the [MASK] token alone.

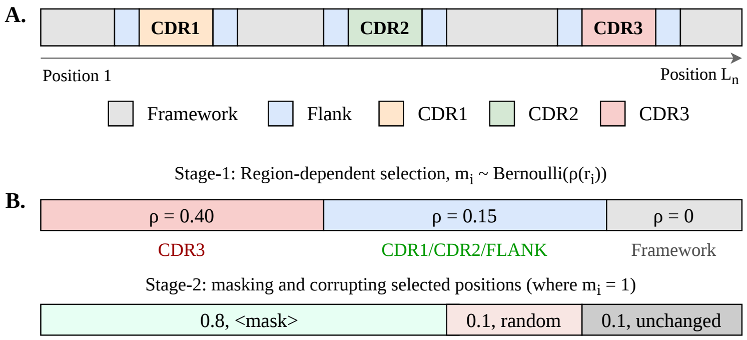


**Figure 1:** Region-aware masking scheme. (a) Schematic antibody sequence with CDR1–3, their four-residue flanks, and framework regions labelled by the priority CDR3 ≻ CDR1 ≻ CDR2 ≻ FLANK ≻ NONEdescribed in Data. (b) The two-stage stochastic corruption process: region-dependent Bernoulli selection (Eq. 10) followed by 80/10/10 corruption of selected positions (Eq. 11); the true residue is retained as the training label in all three branches.

Under a joint length budget $L^{\max}$, the antigen was cropped and the antibody was retained in full. The available antigen budget is

$$B = \max\bigl(1,\ L^{\max} - L_a - 1\bigr), \tag{S3}$$

where the subtracted unit accounts for the separator token, and the antigen was reduced to a contiguous window of width $\min(L_g, B)$, placed at a uniformly random offset during training and centered at evaluation. Because $B$ depends on $L_a$, the effective antigen context varied across examples.

#### 5. LangAAI probability ranking antibody variants

Supplementary Figure 2 repeats the main-text affinity analysis (Figure **??**) for the single-mutant regime. The rank association between joint log-probability and measured affinity persists in both data sets ($\rho = +0.214$ and $+0.216$), and mean affinity gain increases monotonically across score quintiles (Supplementary Figure 2b). A permutation test confirms that neither correlation is an artifact of the marginal distributions of scores or affinities. Shuffling scores against affinities (2,000 permutations) placed the observed $\rho$ at $z = +16$ and $z = +10$ relative to the resulting null (Supplementary Figure 2c). The single-mutant signal is therefore real but weak, and we treat the double-mutant results in the main text as the operative evidence for probability-based ranking.

Supplementary Table 4 reports the same ranking in the form a screening campaign would use – the reliable improver rate within the top-ranked X% of the double-mutant pool. Both campaigns are enriched at every depth examined, but they enrich differently. CoV2-RBD concentrates monotonically as

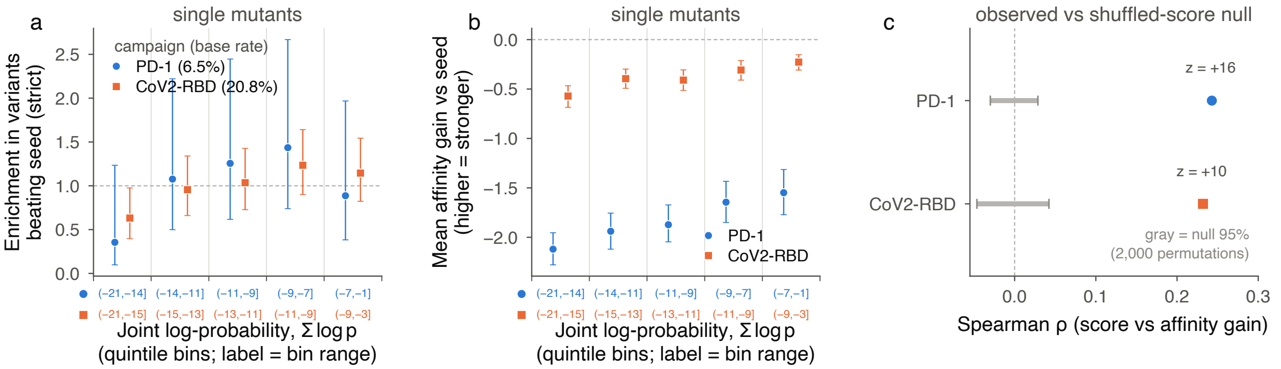


**Figure 2:** The single-mutant regime and the per-position top-k view. **a**, Enrichment in variants beating the seed per quintile of log-probability, single mutants (PD-1 n = 432, CoV2-RBD n = 607; 86–122 variants per quintile). Error bars, 95% Wilson score intervals. **b**, Mean affinity gain per quintile for the same bins; error bars, 95% percentile bootstrap intervals (2,000 resamples). The rank association persists ($\rho = +0.214$ and $+0.216$; P= $7.5 \times 10^{-6}$ and $7.8 \times 10^{-8}$, two-sided Spearman) but the response is shallower than for double mutants: a single substitution spans a narrower joint-probability range, and an order of magnitude fewer variants were measured. **c**, Enrichment over the base rate when proposing the top-k highest-probability residues at every position independently. PD-1 does not exceed its base rate at any k.

the selected fraction narrows, rising from a 12.1% base rate to 29.7% in the top 2% (2.46×). PD-1 peaks early, at 13.5% in the top 20% (1.69×), and does not improve further with more aggressive selection (11.2–12.8% across the top 2–10%). The practical reading is that deeper selection is not uniformly better. For CoV2-RBD, screening the top 2% would test 37 variants to recover 11 reliable improvers, and for PD-1 the same depth yields 11 improvers from 89 variants.

Table 4: Screening precision at decreasing top-ranked fractions of the double-mutant pool. For each campaign, double mutants were ranked by joint log-probability $\sum log p$ and the reliable-improver rate computed within the top X% of that ranking. 100% is the whole population, i.e. the base rate, shown for reference (enrichment = 1.00× by definition). Wilson 95% CI.

| Campaign | Top % | *n* | Reliable improvers | Rate (95% CI) | Enrichment |
|---|---|---|---|---|---|
| PD-1 | 100% (base rate) | 4,460 | 354 | 7.9% (7.2–8.8%) | 1.00× |
| PD-1 | 20% | 892 | 120 | 13.5% (11.4–15.8%) | 1.69× |
| PD-1 | 10% | 446 | 57 | 12.8% (10.0–16.2%) | 1.61× |
| PD-1 | 5% | 223 | 25 | 11.2% (7.7–16.0%) | 1.41× |
| PD-1 | 2% | 89 | 11 | 12.4% (7.0–20.8%) | 1.56× |
| CoV2-RBD | 100% (base rate) | 1,870 | 226 | 12.1% (10.7–13.6%) | 1.00× |
| CoV2-RBD | 20% | 374 | 60 | 16.0% (12.7–20.1%) | 1.33× |
| CoV2-RBD | 10% | 187 | 38 | 20.3% (15.2–26.7%) | 1.68× |
| CoV2-RBD | 5% | 94 | 20 | 21.3% (14.2–30.6%) | 1.76× |
| CoV2-RBD | 2% | 37 | 11 | 29.7% (17.5–45.8%) | 2.46× |

#### 6. Attention weight → NLL correlation

Supplementary Table 5 reports, per joint-stack layer and antibody region, the Pearson correlation between the shift in attention mass directed at the antigen and the change in reconstruction loss under the antigen swap. CDR3 is strongest in every layer (mean $r = 0.250$), roughly three-fold above CDR1 (0.055) and CDR2 (0.083). The framework flank is intermediate (0.169) despite the smallest loss response of any region, so the correlation identifies CDR3 as distinctive rather than tracking the loss hierarchy region-wise.

Table 5: Pearson correlation per layer and CDR region.

| Layer | CDR1 | CDR2 | CDR3 | Flank | Mean |
|---|---|---|---|---|---|
| 0 | 0.0553 | **0.1491** | 0.2499 | 0.1455 | 0.1500 |
| 1 | **0.0767** | 0.0852 | 0.2340 | 0.1855 | 0.1453 |
| 2 | 0.0318 | 0.0538 | 0.2345 | **0.1925** | 0.1282 |
| 3 | 0.0563 | 0.0424 | **0.2800** | 0.1515 | 0.1325 |
| Mean | 0.0550 | 0.0826 | 0.2496 | 0.1687 | – |
